# Network-based functional prediction augments genetic association to predict candidate genes for histamine hypersensitivity in mice

**DOI:** 10.1101/719591

**Authors:** Anna L. Tyler, Abbas Raza, Dimitry N. Krementsov, Laure K. Case, Rui Huang, Runlin Z. Ma, Elizabeth P. Blankenhorn, Cory Teuscher, J. Matthew Mahoney

**Author notes:** Address for correspondence: University Of Vermont Larner College of Medicine, 95 Carrigan Drive, Stafford 118, Burlington, VT 05401.

## Abstract

Genetic mapping is a primary tool of genetics in model organisms; however, many quantitative trait loci (QTL) contain tens or hundreds of positional candidate genes. Prioritizing these genes for validation is often *ad hoc* and biased by previous findings. Here we present a technique for computationally prioritizing positional candidates based on computationally inferred gene function. Our method uses machine learning with functional genomic networks, whose links encode functional associations among genes, to identify network-based signatures of functional association to a trait of interest. We demonstrate the method by functionally ranking positional candidates in a large locus on mouse Chr 6 (45.9 Mb to 127.8 Mb) associated with histamine hypersensitivity (Hhs). Hhs is characterized by systemic vascular leakage and edema in response to histamine challenge, which can lead to multiple organ failure and death. Although Hhs risk is strongly influenced by genetics, little is known about its underlying molecular or genetic causes, due to genetic and physiological complexity of the trait. To dissect this complexity, we ranked genes in the *Hhs* locus by predicting functional association with multiple Hhs-related processes. We integrated these predictions with new single nucleotide polymorphism (SNP) association data derived from a survey of 23 inbred mouse strains and congenic mapping data. The top-ranked genes included *Cxcl12, Ret, Cacna1c*, and *Cntn3*, all of which had strong functional associations and were proximal to SNPs segregating with Hhs. These results demonstrate the power of network-based computational methods to nominate highly plausible quantitative trait genes even in highly challenging cases involving large QTLs and extreme trait complexity.

## INTRODUCTION

Identifying causal variants within quantitative trait loci (QTLs) is a central problem of genetics, but genetic linkage often prevents narrowing QTLs to less than several megabases (Mb). Thus, QTLs may contain hundreds of candidate genes. Instead of revealing the exact gene (or genes) responsible for trait variation, QTL mapping produces positional candidate genes. Rigorously narrowing a QTL by fine mapping with congenic strains can take years or decades, particularly in organisms like mice that have long generation times. Moreover, high-resolution congenic mapping often reveals that the overall QTL effect is due to multiple linked genes within the QTL rather than a single gene (Parker *et al.* 2013; Yazbek *et al.* 2011). Thus, positional data alone are generally insufficient to nominate candidate genes for subsequent biological follow up. To overcome the limitations of mapping data, researchers look within a QTL for plausible candidate genes. However, these selections are typically done by *ad hoc* criteria using prior knowledge or a literature search. This strategy is strongly biased toward prior knowledge and is highly error prone due to missing annotations. There is a need for rigorous and systematic strategies to distinguish among positional candidate genes for mechanistic follow up.

We developed a novel approach to rank positional candidates based on functional association with a trait. To avoid annotation or literature bias, we use functional genomic networks (FGNs), which encode predicted functional associations among all genes in the genome. FGNs such as the Functional Networks of Tissues in Mouse (FNTM) (Goya *et al.* 2015) and HumanBase (Greene *et al*. 2015), are Bayesian integration networks that combine gene coexpression, protein-protein binding data, ontology annotation and other data to predict functional associations among genes. With these networks we can expand on known gene-trait associations to identify sub-networks of trait-associated genes that include novel genes, including in the QTL of interest.

Recent studies with functional genomic networks, for example FNTM, have demonstrated their power to associate novel genes with specific phenotype terms (Guan *et al.* 2010) or biological processes (Ju *et al.* 2013). For example, Guan *et al*. (2010) used a support vector machine (SVM) classifier to identify a gene network associated with bone mineralization and made validated predictions of novel genes that lay outside of all published QTLs for bone mineralization phenotypes (Guan *et al.* 2010). Subsequent studies using similar network-based techniques have made novel predictions of hypertension- and autism-associated genes (Greene *et al.* 2015; Krishnan *et al.* 2016). We have expanded these methods to rank genes in a mapped QTL based on multiple putative functional terms and to integrate these rankings with genetic association *p* values from strain surveys. Our method produces a final ranked list for all genes in the QTL that incorporates both the functional and positional scores of each candidate gene.

Our strategy first builds trait-associated gene lists from structured biological ontologies (e.g., the Gene Ontology (Ashburner *et al.* 2000; Gene Ontology Consortium 2018) and the Mammalian Phenotype Ontology (Smith and Eppig 2012)) and public transcriptomic data from the Gene expression Omnibus (GEO) (Edgar *et al*. 2002; Barrett *et al.* 2012). We then applied machine learning classifiers to the functional networks of tissues in mice (FNTM) (Goya *et al.* 2015) to identify network-based signatures of the trait-related gene lists. This strategy allows us to predict gene-trait associations that are not currently annotated within a structured ontology, overcoming the missing annotation problem.

We applied our approach to a large QTL associated with histamine hypersensitivity (Hhs) in mice. Hhs in mice is a lethal response to a histamine injection. In insensitive mice, a histamine injection produces an inflammatory response that resolves without further treatment. Mice with the Hhs response develop excitation and ear blanching, followed by progressive respiratory distress, vasodilation, anaphylactic shock, and death (Vaz *et al.* 1977; Wang *et al.* 2014). Hhs can be induced in a subset of mouse strains by sensitization with Complete Freund’s Adjuvant (CFA). Hhs also develops spontaneously in SJL/J animals older than six months of age.

We previously mapped Hhs to a locus on Chr 6 (45.9 Mb to 127.8 Mb; the *Hhs* locus), which was confirmed using a congenic line (B10.S-*Hhs*^*SJL*^) (Raza *et al.* Under Review). Because of the large size of this locus, additional information is required to identify causal variants. To narrow down candidates, we integrated novel genetic association data from interval-specific congenic recombinant lines (ISCRLs) and an inbred strain survey with our network-based functional predictions of Hhs-related genes. By augmenting positional data with functional predictions, we dramatically reduced the candidate gene list to a tractable set of high-quality candidates that are implicated in Hhs-related processes.

## MATERIALS AND METHODS

As a supplement to the methods section, this paper includes an executable workflow (Figure 4). For additional details about specific parameters and inputs, please see the workflow (Figure 4, Data Availability; also available at https://github.com/MahoneyLab/HhsFunctionalRankings).

**Figure 1.**
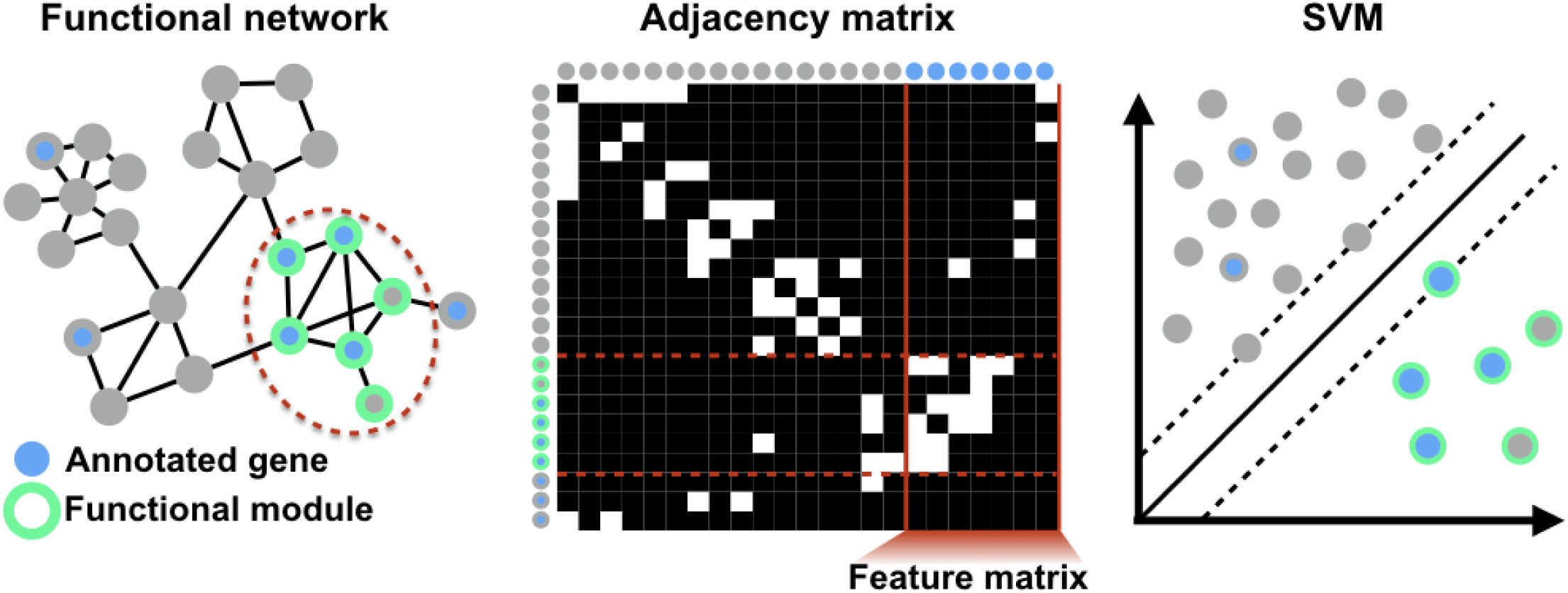
Network-based machine learning for functionally annotating genes. **A** Known-positive genes annotated to a functional term (blue nodes) are typically densely interconnected in a functional network. **B** The adjacency matrix of a network is a tabular representation of the connectivity structure of the network in which each row/column corresponds to a node of the network, and connected pairs of nodes have non-zero values in the corresponding cell of the matrix. Note that in general the connections are weighted, but for display we are only showing present/absent links (white/black cells). The connections from every gene in the genome to the known positives form a sub-matrix of the adjacency matrix called the feature matrix (vertical red lines), whose rows are the feature vectors for each gene. **C** Using the network-based feature vectors for each gene, we train SVMs to distinguish known positives (blue dots) from random genes in the genome (gray dots) to identify the full sub-network corresponding to the true positive genes (green bordered dots and dotted red lines in panels **A**,**B**).

**Figure 2.**
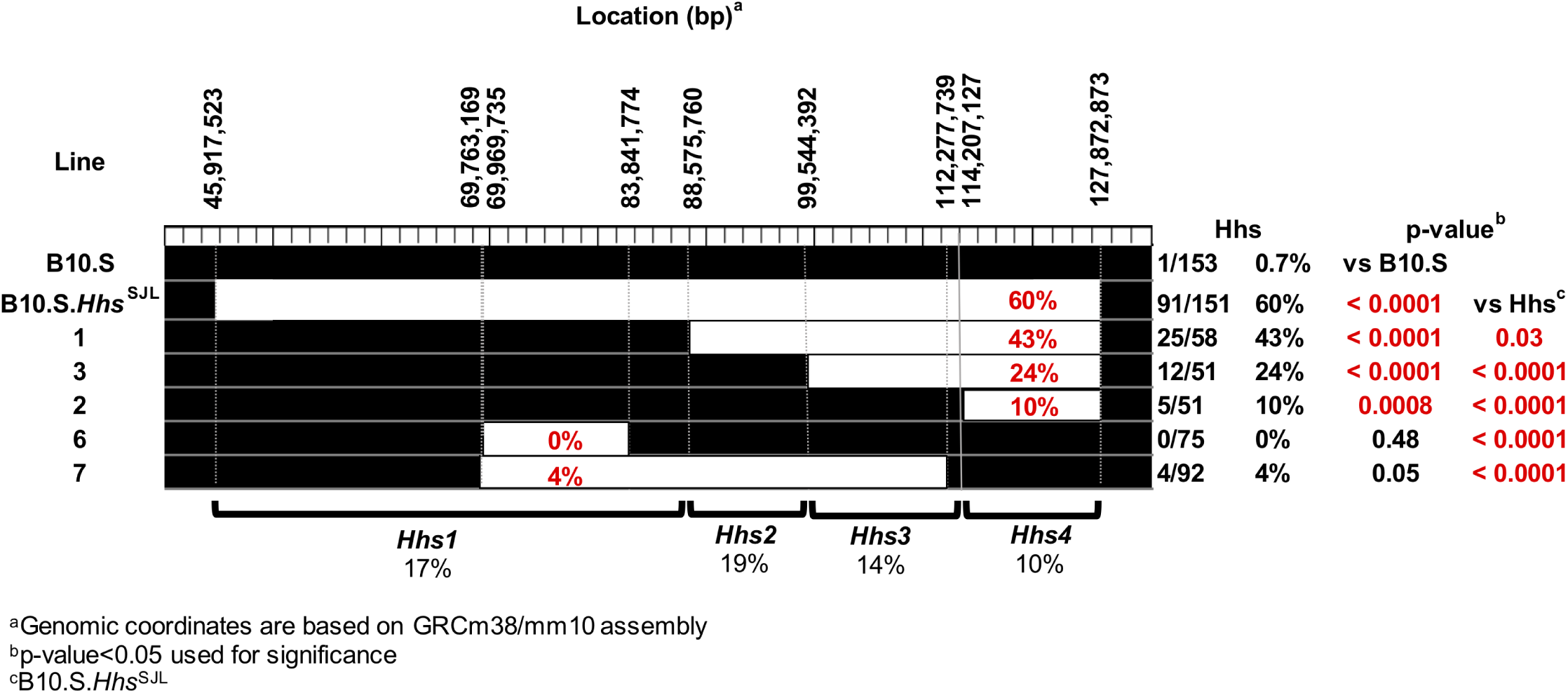
Interval specific recombinant congenic (ISRC) mapping places Hhs candidates in four genetic loci. ISRC lines were injected (D0) with complete Freund’s adjuvant (CFA) and subsequently challenged (D30) with and i.v. injection of histamine to determine histamine hypersensitivity. Deaths were recorded at 30 min post injection and the data are reported as the number of animals dead over the number of animals studied. Significance of observed differences was determined by a *χ*^2^ test with *p*-values <0.05 considered significant.

**Figure 3.**
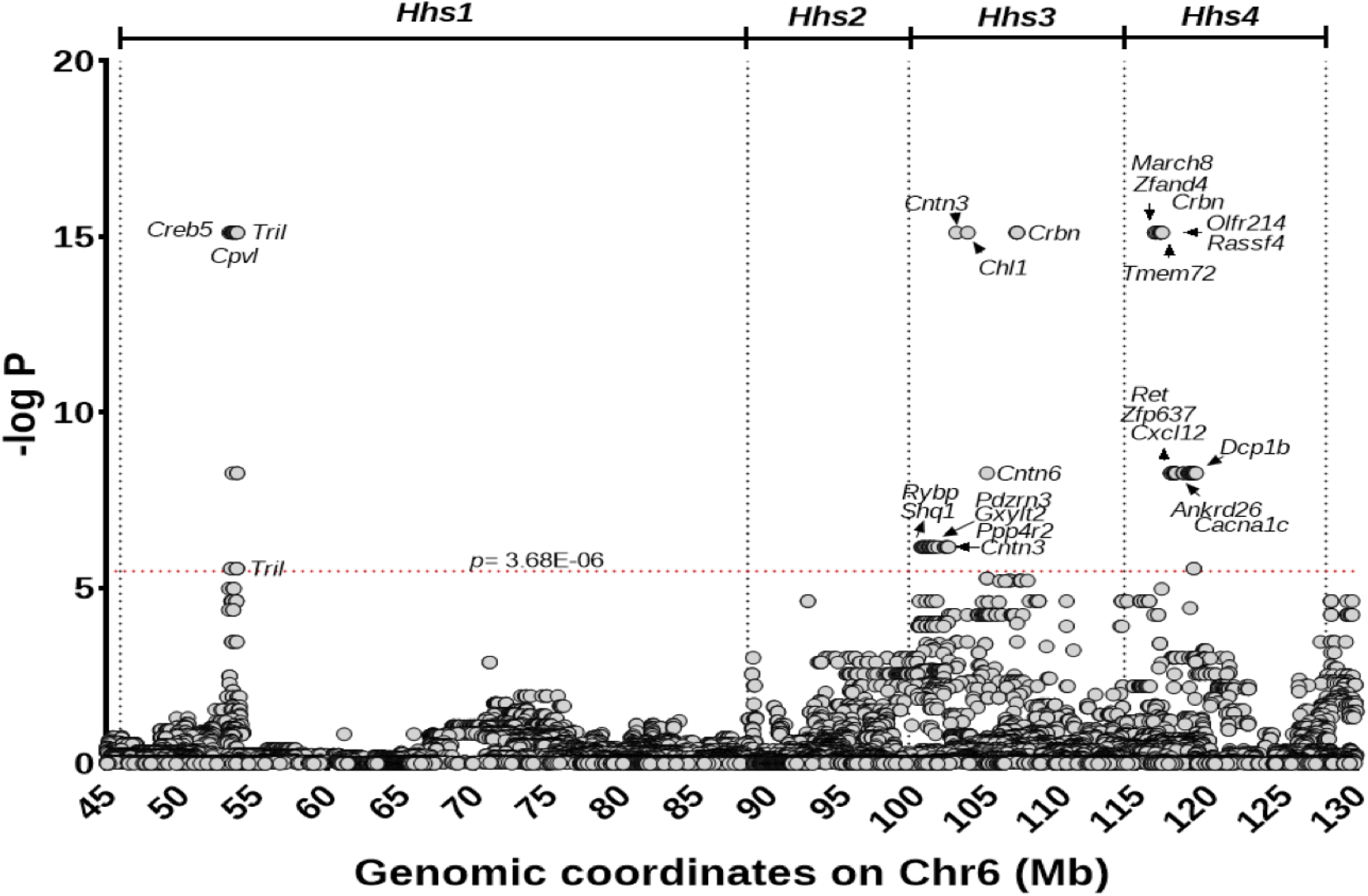
Targeted genetic association analysis for Hhs. Negative log-transformed *p* values of SNP associations with Hhs. Genomic coordinates (mm10 Mbp) of each SNP are shown along the *x*-axis. Each circle denotes a single SNP. Gene names are included for SNPs that crossed *p*-value threshold of 3.68 × 10^*-*6^ shown with a red dotted line. The location of Hhs sub-QTLs are shown at the top of the figure.

**Figure 4.**
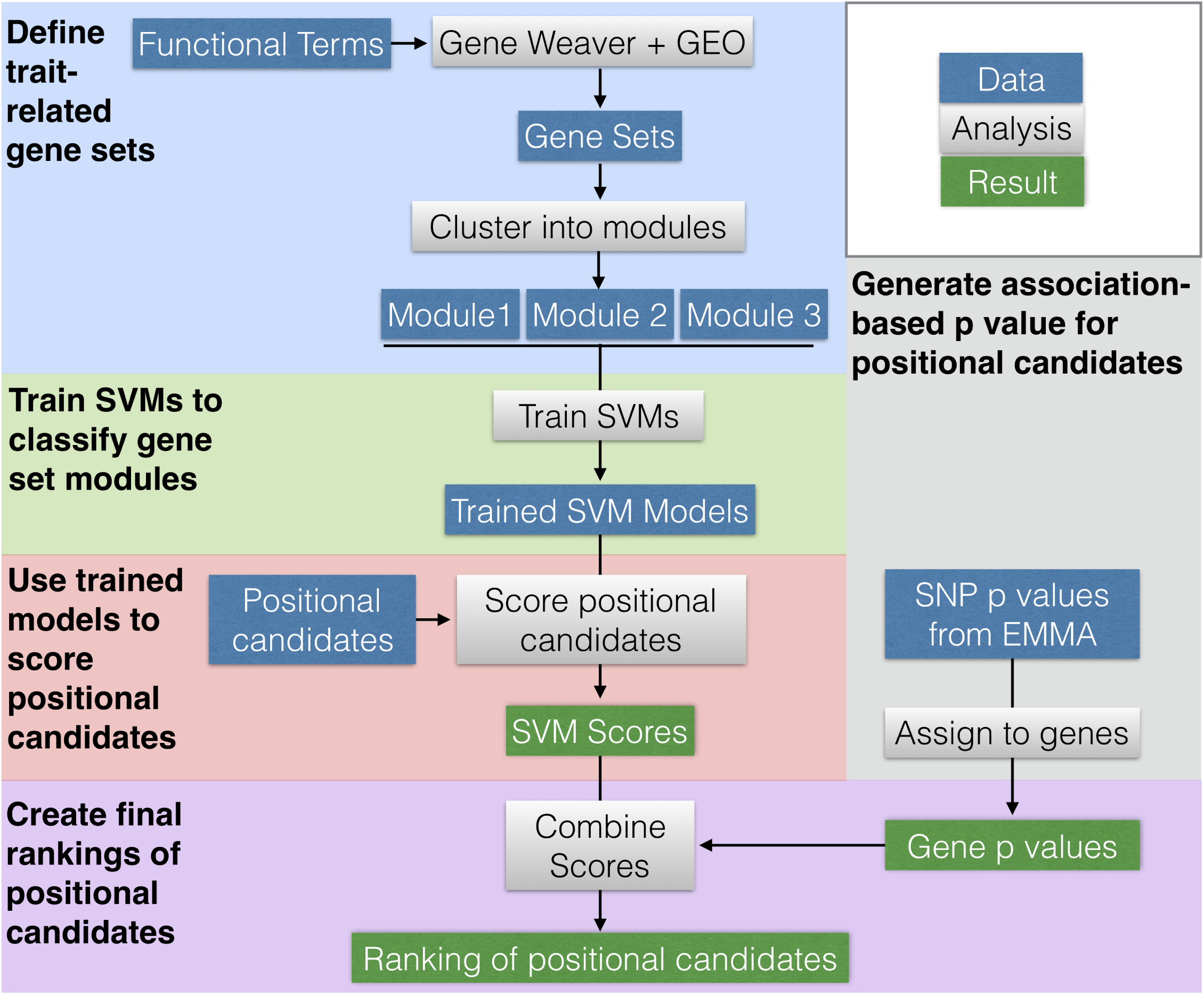
Workflow Overview. The workflow is broken into blocks by color, each with a bolded title. Each block shows how data (blue rectangles) were operated on (gray rectangles) to achieve results (green rectangles). Arrows show the general flow of work and dependence (and independence) of individual analyses.

### Animals

A total of 23 mouse strains (129X1/SvJ, A/J, AKR/J, B10.S-*H*2s/SgMcdJ (B10.S), BALB/cJ, BPL/1J, BPN/3J, C3H/HeJ, C57BL/6J, CBA/J, CZECHII/EiJ, DBA/1J, DBA/2J, FVB/NJ, JF1/MsJ, MOLF/EiJ, MRL/MpJ, MSM/MsJ, NOD/ShiLtJ, NU/J, PWD/PhJ, PWK/PhJ, SJL/J and SWR/J were purchased from the Jackson Laboratory (Bar Harbor, ME). All mice, including B10.S-HhsSJL and B10.S-HhsSJL ISRC lines, were generated and maintained under specific pathogen-free conditions in the vivarium of the Given Medical Building at the University of Vermont according to National Institutes of Health guidelines. All animal studies were approved by the Institutional Animal Care and Use Committee of the University of Vermont.

### Hhs Phenotyping

On day (D) 0 mice were injected i.p. with complete Freund’s adjuvant (CFA) (Sigma-Aldrich, St. Louis, MO) supplemented with 200 *µ*g of *Mycobacterium tuberculosis* H37Ra (Difco Laboratories, Detroit, MI). On D30 histamine hypersensitivity was determined by i.v. injection of histamine (mg/kg dry weight free base) in phosphate buffered saline (PBS). Deaths were recorded at 30 min post injection and the data are reported as the number of animals dead over the number of animals studied. Significance of observed differences was determined by Chi-square with *p*-values <0.05 significant.

### DNA extraction and genotyping

DNA was isolated from mouse tail clippings as previously described (Sudweeks *et al.* 1993). Briefly, individual tail clippings were incubated with 300*µ*L cell lysis buffer (125*µ* g/mL proteinase K, 100 mM NaCl, 10mM Tris-HCl (pH 8.3), 10 mM EDTA, 100mM KCl, 0.50% SDS) overnight at 55°C. The next day, 150*µ*L of 6M NaCl were added followed by centrifugation for 10 min at 4°C. The supernatant layer was transferred to a fresh tube containing 300*µ*L of isopropanol. After centrifuging for two minutes, the supernatant was discarded, and pellet washed with 70% ethanol. After a final two min centrifugation, the supernatant was discarded, and DNA was air dried and resuspended in 50*µ*L TE.

#### Genotyping

Genotyping was performed using either microsatellite markers in a standard PCR reaction or sequence specific SNP primers in a phototyping reaction. Polymorphic microsatellites were selected to have a minimum polymorphism of 8bp for optimal identification by agarose gel electrophoresis. Briefly, primers were synthesized by IDT-DNA (Coralville, IA) and diluted to a concentration of 10*µ*M. PCR amplification was performed using Promega GoTaq. The cycling conditions included a two-minute initial denaturation step at 94°C followed by 35 cycles of 94°C for 30 seconds, 55°C for 30 seconds and 72°C for 30 seconds followed by a final extension step at 72°C for five minutes. Amplicons were subjected to 2% agarose gel electrophoresis and visualized by ethidium bromide and UV light.

#### Phototyping

Genotyping was performed using sequence-specific primers that differ only at the 3’ nucleotide corresponding to each allele of the identified SNP (Bunce *et al.* 1995). Each primer set was designed using Primer3 to have a Tm of 58-60°C, synthesized by IDT-DNA (Coralville, IA), and used at a concentration of 100*µ*M (primer sequences are available in Supplemental File 1). PCR reactions were subjected to multistage (high, medium and low stringency) cycling conditions as described and if found to be necessary, the cycle conditions at each stage were adjusted to accommodate the optimal annealing temperature. Amplicons were electrophoresed with 10*µ*L Orange G loading buffer on a 1.5% agarose gel stained with ethidium bromide and visualized by UV light. The presence of a SNP specific allele was scored by observing an amplicon of the expected size in either reaction. Cycling conditions are available in Supplemental File 6.

### Generation of Hhs congenic lines and GigaMUGA

B10.S-*Hhs*^*SJL*^ ISRC lines were generated by identifying recombinant haplotypes across the Hhs interval among (B10.S-*Hhs*^*SJL*^ × B10.S) × B10.S BC1 mice and then fixed as homozygous lines (Figure 2). To identify potential contaminating background loci segregating among the strains and to further refine the recombination break points of each line, the lines were further genotyped using GigaMUGA arrays (143,259 markers) by the commercial service of Neogen/Geneseek (Lincoln, NE).

### Targeted genetic association testing

We retrieved genotype data (both coding and non-coding) of the 23 mouse strains from the databases at the Sanger Institute (https://www.sanger.ac.uk/science/data/mouse-genomes-project) and The Jackson Laboratory (https://phenome.jax.org/). The lack of representation of wild-derived strains e.g., MOLF and others, in these databases were compensated by genotyping using high-throughout Nimblegen sequence capture (^®^SeqCap EZ Target Enrichment www.sequencing.roche.com). All these data sources were collated to generate genotype information for a total of 13,598 SNPs across the *Hhs* locus (45-128 Mbp, Additional File 8). To calculate associations between genetic polymorphisms and Hhs, we used efficient mixed-model association (EMMA) (Kang *et al.* 2008). This method treats genetic relatedness as a random variable in a linear mixed model to account for population structure, thereby reducing false associations between SNPs and the measured trait. We used the likelihood ratio test function (emma.ML.LRT) to generate *p* values. Significance was defined with a Bonferroni correction (*p* = 0.05/13, 598). Genomic coordinates included for each SNP using the latest mouse genome build GRCm38.p5/mm10.

### Trait-related gene sets

The positional candidate genes were ranked based on their predicted association with seven functional terms related to the Hhs phenotype: “aging”, “mycobacterium tuberculosis”, “cardiac”, “G-protein coupled receptor”, “histamine”, “inflammation”, “type I hypersensitivity”, and “vascular permeability.” We used Gene Weaver (Baker *et al.* 2012) to identify genes associated with each term. We entered each term into the Gene Weaver homepage (https://geneweaver.org). We restricted the search to human, rat, and mouse genes, and to curated lists only. Mouse homologs for each gene were retrieved using batch query tool in MGI (http://www.informatics.jax.org/batch_data.shtml). In addition, we used Gene Expression Omnibus (GEO) and PubMed to retrieve expression data sets for each phenotype term. The data sets used are listed in Supplemental File 7. Final gene lists consisted of the unique set of genes associated with each process term.

### FNTM network

We trained support vector machines (SVMs) to classify genes in each gene list using features derived from the Functional Network of Tissues in Mouse (FNTM) (Goya *et al.* 2015). In this network, genes are nodes, and the edge weights between them are continuous values between 0 and 1 predicting the degree to which each pair of genes is functionally related. Larger values indicate higher predicted functional relatedness. Functional relatedness in this network was predicted through Bayesian integration of data sets from multiple sources, including gene expression, protein-protein interaction data, and annotation to GO terms (Goya *et al.* 2015). We downloaded the top edges of the mouse network on January 15, 2018 from https://http://fntm.princeton.edu.

### Clustering gene sets

Guan *et al*. (2010) noted that support vector machines trained on 200 to 300 genes yielded the best classification accuracy. Two of our gene lists had fewer than 100 genes. For all lists over 400 genes, we reduced the size of our training sets by clustering each term gene set into modules using the fast greedy (Newman 2004) algorithm in the R/igraph package (Csardi 2006). We applied the fast greedy algorithm iteratively until all modules comprised fewer than 400 genes (Supplemental Table 2). Using a maximum modules size of 300 overly fragmented the networks yielding many modules with fewer than 100 genes.

### Machine learning

To classify novel genes as belonging to a functional module, we trained SVMs using the connection weights in the FNTM network as features, as described in Guan *et al*. (2010). Briefly, an annotated set of genes (Figure 1A, blue nodes) is used as a set of known positives for the corresponding functional module. Other genes in this module are expected to be strongly functionally connected to these known positives, i.e. have high probability of functionally interacting with known positives. Each gene, therefore, is represented as a *feature vector* of connection weights to the known positives, which can be visualized as a sub-matrix of the *adjacency matrix* of the network (Figure 1B). Correspondingly, the rows of this matrix are labeled as either known positive or not (Figure 1B, blue dots vs. gray dots). We used the e1071 package in R (Meyer *et al.* 2018) to train SVMs to distinguish the known positive genes from an equal-sized set of genes selected at random from outside the known positive list using the network-based feature vectors (Figure 1C). The trained model can then annotate novel genes as belonging to the functional module by classifying all gene in the genome (Figure 1C, green bordered nodes).

We trained 100 SVMs on each module selecting a new set of random genes for each run. We used a linear kernel and 10-fold cross-validation for each SVM. We trained each SVM over a series of cost parameters. We started with the sequence 1 × 10^*-*5^ to 1 × 10^2^ by factors of 10, and iteratively narrowed the range of cost parameters until we found a series of eight cost parameters that maximized the accuracy of the SVM (see Workflow).

We calculated the area under the ROC curves (AUC) over all runs in the following way: For a sequence ranging from the minimum SVM score to the maximum SVM score, we quantified all true positives (TP), true negative (TN), false positives (FP) and false negatives (FN). The TP genes in this case were those genes from the known positives that were correctly classified as being in the module (above the SVM score cutoff). TN genes in this case were those genes outside the module that were correctly classified as being outside the module (below the SVM score cutoff). We calculated the AUC across the average curve for all 100 SVMs for each module.

### Positional Candidate Scoring

We used the trained SVMs to score each positional candidate gene in the *Hhs* locus. The score for each gene gave an estimate of how functionally related each gene was to each module based on its connection weights to the known module genes in the FNTM mouse network. Genes with large positive scores were predicted by the SVMs to interact functionally with the genes in the module, while genes with negative scores were predicted to *not* functionally interact with the module genes. To be able to compare SVM scores across different trained models, we calculated a false positive rate (*FPR*) for each gene and each SVM as follows: For each gene’s SVM score we calculated the number of true positives (*TP*), true negatives (*TN*), false positives (*FP*) and false negatives (*FN*) classified by the SVM. The *FPR* for a given SVM score was calculated as *FP*/(*FP* + *TN*).

The final functional score for each was the *max*(*-log*_10_ (*FPR*)) across all modules. This meant that genes with a high functional score for a single module, but low functional scores for other modules received higher overall scores than genes with moderately high scores across all modules.

### Combined Gene Score

High-quality candidate genes in the locus should not only be functionally related to the trait of interest, but should also segregate with the trait of interest. We thus defined a combined gene score (*S*_*cg*_) that combined these two aspects of the analysis:

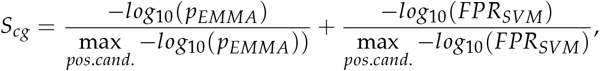

where the denominators of the two terms on the right hand side are the maximum values of *-log*_10_ (*p*_*EMMA*_) and *-log*_10_ (*FPR*_*SVM*_) over all positional candidates in *Hhs*, respectively, which normalizes the functional and positional scores to be comparable to each other. EMMA *p* values for SNPs were assigned to the nearest gene within 1 megabase using the R package biomaRt (Durinck *et al*. 2005, 2009) (Supplemental Table 3). Genes for which more than one SNP was assigned were given the maximum *-log*_10_ (*p*_*EMMA*_) across all SNPs associated with that gene. The full matrix of combined scores across all gene sets is in Supplemental Table 5. The rows of this matrix are sorted by the maximum gene score across all gene lists.

## RESULTS

### Generation of Interval Specific Recombinant Congenic Lines (IS-RCL) across the *Hhs* locus

In prior work, we mapped the genetic locus regulating susceptibility to age- and/or inflammation (CFA)-dependent sensitivity to histamine on Chr 6 in SJL mice (Raza *et al.* Under Review). The B10.S-*Hhs*^*SJL*^ congenic mice exhibit Hhs and carry a large ≈ 83 Mb region of SJL between 45.9 Mb to 127.8 Mb on the resistant B10.S background (Raza *et al.* Under Review). This large QTL includes 628 protein coding genes. To narrow this region, we generated five ISRCLs using B10.S-*Hhs*^*SJL*^ x B10.S backcross mice and assessed their susceptibility to Hhs (Figure 2). Under an additive model, these data suggest that *Hhs* is composed of four sub-QTL which we have designated *Hhs1, Hhs2, Hhs3*, and *Hhs4*, each contributing 17%, 19%, 14% and 10%, respectively, to the overall penetrance. Importantly, for each sub-QTL this makes positional candidate gene identification using interactive high resolution congenic mapping impractical.

### Inbred strain survey of Hhs

To investigate whether the Hhs phenotype is unique to SJL, we assessed histamine responses for 23 inbred mouse strains (including SJL and B10.S; Table 1). These strains were chosen using haplotype structure across the Hhs interval to identify additional mouse strains that are likely to share a susceptible *Hhs* allele (data not shown). 129X1/SvJ, ALR/LtJ, BPN/3J, FVB/NJ, NOD/ShiLtJ, NU/J, SJL/BmJ and SWR/J mice were identified as having similar haplotype structure as SJL at the *Hhs* locus. ALR/LtJ and SJL/BmJ mice required embryo recovery and were therefore not included. Hhs phenotyping identified FVB/NJ, SWR/J, and NU/J mice as Hhs-susceptible, whereas 129/X1/SvJ, NOD/ShiLtJ, and BPN/3J were resistant. Taken together with our earlier data, these results indicate that Hhs susceptibility segregates among a unique subset of SJL/J-related strains (Petkov *et al.* 2004).

**Table 1.** A survey Hhs phenotypes across 23 inbred mouse strains.

| Strain | HA | Strain | HA | Strain | HA | Strain | HA |
| --- | --- | --- | --- | --- | --- | --- | --- |
| 129X1/SvJ | 0/8 | C3H/HeJ | 0/8 | DBA/2J | 0/8 | PWK/PhJ | 0/6 |
| A/J | 0/8 | C57BL/10J | 0/8 | JF1/Ms | 0/8 |  |  |
| AKR/J | 0/8 | C57BL/6J | 0/7 | MOLF/EiJ | 0/8 | FVB/J | 6/8 |
| BALB/cJ | 0/8 | CBA/J | 0/8 | MRL/MpJ | 0/8 | NU/J | 5/8 |
| BPL/1J | 0/8 | CZECHII/EiJ | 0/8 | NOD/ShiLtJ | 0/8 | SJL/J | 12/12 |
| BPN/3J | 0/8 | DBA/1J | 0/8 | PWD/PhJ | 0/12 | SWR/J | 6/8 |

Cohorts of CFA injected 8- to 10-week old mice were challenge 30 days later with 75 mg/kg HA by i.v. injection, and deaths recorded at 30min. Results are expressed as the (number of animals dead)/(number of animals studied).

### Targeted genetic association analysis for *Hhs*

Our result from previous linkage analysis (Raza *et al.* Under Review) and congenic mapping localized *Hhs* to an ≈ 83 Mb region on Chr 6 between 45.9 Mb to 127.8 Mb. Given that Hhs-susceptibility is restricted to a unique subset of inbred strains, particularly the closely related SJL/J, FVB/NJ, and SWR/J, we performed a targeted association analysis between SNPs in the *Hhs* locus across all 23 inbred strains (*cf*. Benson *et al*. (2017)).

We tested the association of 13,598 SNPs across the *Hhs* locus using efficient mixed-model association (EMMA) (Kang *et al.* 2008). A total of 84 SNPs in 23 genes showed significant associations (*p* ≤ 3.68 × 10^*-*6^) (Figure 3, Table 4). The majority of the significant hits were intronic (71%), non-coding (12%), intergenic (4%) or regulatory (5%) variants. Interestingly, there was overlap between three of the four *Hhs* sub-QTLs (Figure 2) and SNP-association peaks.

### Network-based prediction of Hhs-associated genes

To predict functional candidates among the positional candidates in the *Hhs* locus, we delineated a list of Hhs-associated biological processes and trained machine learning classifiers to identify sub-networks of functional genomic networks associated with each of these processes. An overview of our workflow is in Figure 4. We first defined gene sets that were related to seven terms that are functionally related to the Hhs phenotype.

The terms and their justifications are as follows:

- *Type I hypersensitivity/Anaphylaxis*: The death response following systemic histamine challenge exhibits symptoms of type I hypersensitivity/anaphylaxis including respiratory distress, vasodilation, and anaphylactic shock (Vaz *et al.* 1977).
- *Cardiac*: There is evidence suggesting that anaphylactic shock in mice is associated with decreased cardiac output, rather than solely a function of systemic vasodilation (Wang *et al*. 2014).
- *Histamine*: Hhs is elicited by a systemic histamine challenge (Raza *et al.* Under Review).
- *G-protein coupled receptor*: Histamine receptor H1 (*Hrh1*) signaling is required for the Hhs phenotype, and all histamine receptors belong to the family of G-protein coupled receptors (Hill *et al.* 1997).
- *Aging*: Spontaneous Hhs develops after six months of age in sensitive mouse strains (Raza *et al.* Under Review).
- *Inflammation*: Treatment with pro-inflammatory CFA induces Hhs in sensitive mouse strains.
- *Tuberculosis*: Hhs is induced in some mouse strains by CFA, which contains inactivated *Mycobacterium tuberculosis* (Raza *et al.* Under Review).
- *Vascular permeability*: The Hhs response includes vascular leakage in skin and skeletal muscles as assessed by Miles’ assay (Raza *et al.* Under Review).

We used Gene Weaver, the Gene Expression Omnibus (GEO), and PubMed to retrieve gene sets associated with each of these terms (see Materials and Methods). The gene sets ranged in size from 651 to 1466 genes. Because Guan *et al*. (2010) found that SVMs trained on gene sets with around 300 genes performed best for network-based functional prediction, we clustered large gene sets into modules of approximately 300 genes and analyzed each module separately (see Materials and Methods). Supplementary Table 2 shows the number of genes in each module, as well as the top five enrichment terms for each using the R package gProfileR (Reimand *et al.* 2018). Multiple members of these gene sets are encoded in the *Hhs* locus. For example, e.g. *Hrh1* was a member of the Anaphylaxis gene set. To reduce bias in classification, we removed all such genes from each gene set before SVM training. We then trained an ensemble of 100 SVMs on each module gene set. We calculated ROC curves for each model to quantify the ability of each set of SVMs to distinguish genes inside the module gene set from all genes outside the module gene set. AUCs ranged from 0.9 to 0.975 indicating that the SVMs were able to classify the genes in each list robustly. In other words, each gene set used to define a putative Hhs-related process forms a distinct subnetwork of the full functional genomic network.

We then applied the trained SVM models to the positional candidate genes in the *Hhs* locus. By classifying each positional candidate, we can identify genes that are likely to be functionally associated with each module gene set. For example, for the Anaphylaxis module gene set, the histamine receptor *Hrh1* received a positive score indicating that the SVMs predicted it belonging to the Anaphylaxis gene set despite its absence from the training set. This example provides a positive control and shows that the SVMs identify biologically relevant patterns in the functional genomic network. In addition to the SVM score, we calculated a false positive rate (*FPR*) for each gene (see Materials and Methods). Low *FPR*s indicate high confidence in the classification. The details of this analysis are described in an executable workflow as a companion to this paper (see Data Availability).

### Integration of functional enrichment with genetic association

Genes that are predicted to be highly functionally related to the trait may not have functionally variant alleles in the study population, and may therefore be unlikely to drive the observed strain differences in Hhs. To identify genes that were likely to have functionally relevant polymorphisms, we integrated functional scores with SNP association *p* values to focus only on those candidates that satisfied both criteria. By plotting the maximum functional score for a gene, *-log*_10_ (*FPR*_*SV M*_) versus the *-log*_10_ (*p*_*EMMA*_) (normalized to the max values; see Materials and Methods), we can identify genes that were predicted to be both highly functionally related to Hhs phenotype and likely to have functional polymorphisms that segregated with Hhs susceptibility (Figure 5). The blue line in Figure 5 traces along the Pareto front of the gene set in this space. For any gene on this line, finding a gene with a stronger functional association means finding a gene with lower SNP *p* value, and *vice versa*. The genes near the Pareto front have either segregating polymorphisms or are predicted to be functionally related to Hhs, or both. All such genes are potentially good candidates for experimental follow-up.

**Figure 5.**
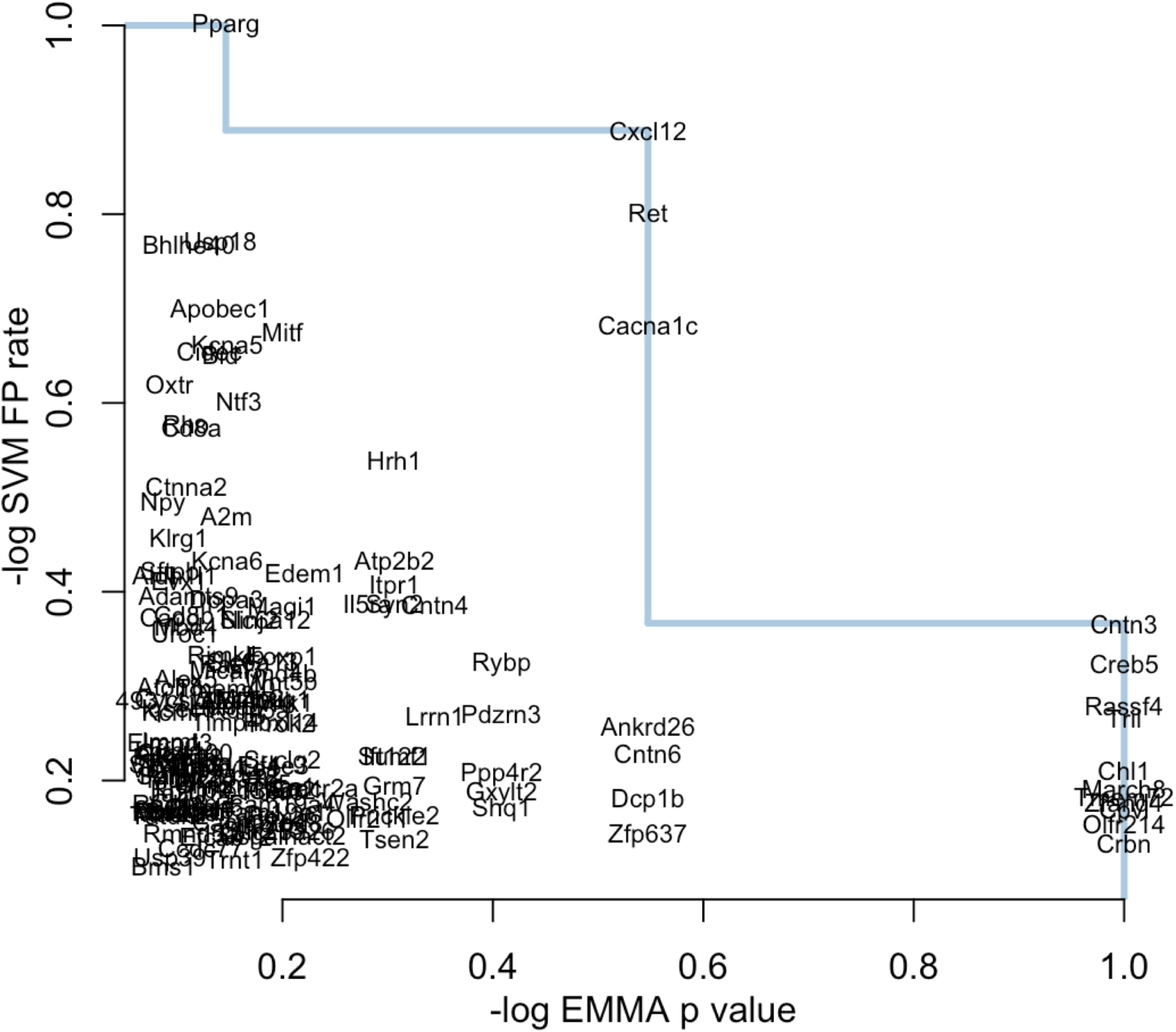
Two axes of gene scoring. Gene names are plotted by their *-log*_10_ (*p*_*EMMA*_) on the *x*-axis and the *-log*_10_ (*FPR*_*SV M*_) on the *y*-axis. Both scores were scaled by their maximum value for better comparison. Genes farther to the right were associated with SNPs that segregated with Hhs. Genes higher up on the *y*-axis are associated with stronger functional association with gene modules. The blue line marks the Pareto front. Genes on this line maximize the two scores and are the best candidates based on the combination of both scores.

To rank the candidates with a single score, we defined a final gene score (*S*_*cg*_) for each gene, which is the sum of the (normalized) *-log*_10_ (*FPR*) and the *-log*_10_ (*p*_*EMMA*_) (Figure 6). This score prioritizes candidates in the upper right quadrant with simultaneously high positional and functional scores. The genes in the upper right quadrant—*Cxcl12, Ret* and *Cacna1c*—had near-maximal scores along both axes and were therefore ranked as the best candidates for follow-up. The full table of gene scores by module can be seen in Table 5.

**Figure 6.**
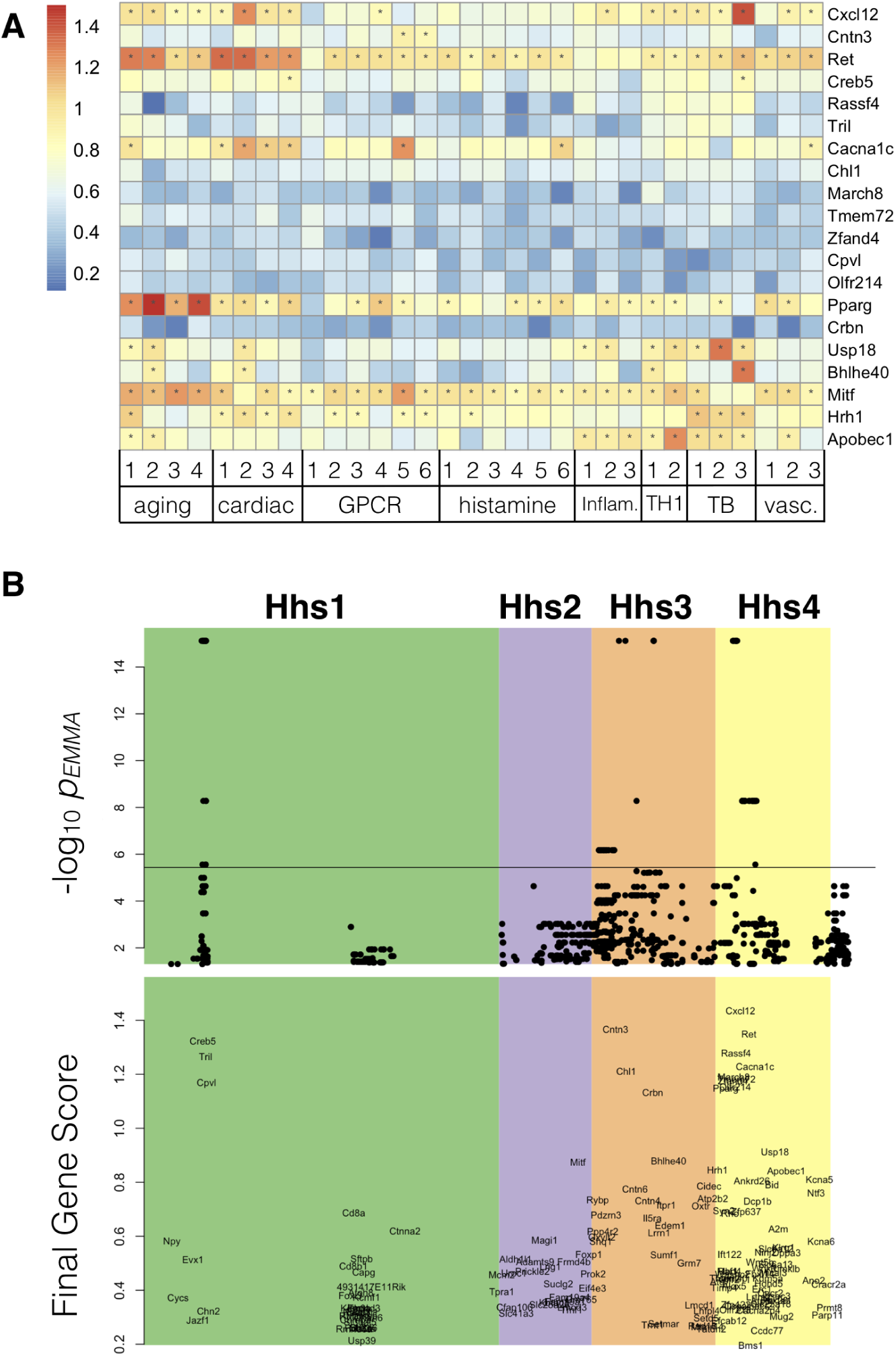
Final gene scores. Gene functional values were combined with SNP associations to assign each gene a final gene score (*S*_*cg*_). Higher gene scores indicate better candidates. **A** Heat map showing the final score of each of the top 20-ranked genes for each gene module. To aid visualization of the strongest candidates, asterisks in each cell indicate where candidate genes were associated with a module with an *FPR*_*SVM*_ ≤ 0.2. **B** The top panel shows individual SNPs plotted at their genomic location (*x*-axis) and their log10 (*p*_*EMMA*_) (*y*-axis). All SNPs with nominally significant p value (*p* ≤ 0.05) are plotted. The horizontal line indicates the Bonferroni corrected significance cutoff (*p* ≤ 0.05/13598). The four sub-QTLs are demarcated by background color and are labeled at the top of the figure. The bottom panel shows genes plotted at their genomic location (*x*-axis) and their final gene score (*S*_*cg*_) (*y*-axis) to demonstrate how the final ranked genes align with the SNP association data.

In addition to identifying the top-ranked gene over the full *Hhs* locus, we identified a top-ranked gene for each sub-QTL identified through congenic mapping. Figure 6A shows the functional associations across all modules of the top 20 genes ordered by final gene score (*S*_*cg*_). The full matrix of scores for all ranked genes can be found in Supplemental Table 5.

## DISCUSSION

In this analysis, we identified a small set of positional candidate genes in a large locus by combining computational predictions of functional association with Hhs and SNP associations. The final list of genes is highly plausible and can be followed up relatively easily with modern genetic editing techniques.

### High-quality candidates for *Hhs*

Three genes in the final ranked list deserve particular attention: *Cxcl12, Ret*, and *Cacna1c*. Of all genes in the locus, these three lie on the Pareto front with both low genetic association *p* values and high functional scores (Figure 5). The top-ranked gene, *Cxcl12* (a.k.a. stromal cell-derived factor 1), is chemotactic for mast cells via the chemokine receptor *Cxcr4* (Ghannadan *et al.* 2002). Mast cells are major drivers of pathological events in anaphylaxis (Lieberman and Garvey 2016), demonstrating that the final predictions are highly relevant to Hhs. The second-ranked gene *Ret* encodes a pleiotropic tyrosine protein kinase involved in cell differentiation, growth, migration, and survival (Motenko *et al.* 2015), inflammation (Rusmini *et al.* 2013) and the development of the cardiovascular system (Hiltunen *et al.* 2000). Alleles of this gene could conceivably modify multiple processes underlying Hhs, including the both the anatomical background susceptible to Hhs and the acute response to histamine. *Ret* was significantly associated with multiple functional gene sets (Figure 6A). The third-ranked gene, *Cacna1c*, encodes the voltage-dependent calcium channel Ca_v_1.2, which is expressed in the heart, muscle, and endocrine glands (Mouse Genome Informatics Mouse Genome Informatics Web Site). Mutations in *Cacna1c* are associated with electrophysiological alterations in the heart (Napolitano *et al.* 2015; Hedley *et al*. 2009) suggesting a possible role for *Cacna1c* in impaired cardiac function in Hhs. Interestingly, SNPs in human *CACNA1C* were recently associated with chronic spontaneous urticaria (i.e., spontaneous episodes of hives and/or angioedema) and antihistamine drug response (Yan *et al.* 2018)(paper in Chinese). These results suggest a direct connection between *Cacna1c* and anaphylactic or hypovolemic shock.

All of the above genes lie in the *Hhs4* locus, which accounts for only a portion of the total variation in the Hhs phenotype. In the *Hhs3* locus, the highest-ranked candidate gene was *Cntn3*, which encodes for contactin 3, an activator protein of the small GTPase *Arf. Cntn3* is a member of the contactin family of immunoglobulins. Genetic variants of human *CNTN3* are associated with an enlargement of the aorta, acute heart rate recovery, and abdominal aortic aneurysm, suggesting a potential connection to impaired cardiac function during histamine challenge (Elmore *et al.* 2009). Intriguingly, *CNTN3* is near a segregating SNP for Systemic Capillary Leak Syndrome (SCLS) from a human GWAS. SCLS is an extremely rare disease characterized by transient but potentially lethal episodes of diffuse vascular leakage of proteins and fluids into peripheral tissues, resulting in massive whole-body edema and hypotensive shock. The pathological mechanisms and genetic basis for SCLS remain elusive (Xie *et al.* 2013), but SCLS shares many phenotypic properties with Hhs in mice. In particular, SCLS attacks are diagnosed based on the clinical triad of hypotension, elevated hematocrit, and hypoalbuminemia, all of which naturally occur in the Hhs-sensitive SJL mouse strain (Raza *et al.* Under Review). The potential association between *CNTN3* and SCLS, therefore, lends credence to its possible functional role in Hhs as well. Indeed, *CNTN3* was not only a positional candidate in the SCLS GWAS, but was contained within functional terms that were enriched among the top positional candidate genes (cf. Table 5 of Xie *et al*. (2013)), indicating that CNTN3 functions in concert with other genetic risk factors for SCLS.

In the *Hhs1* locus, the top hits in were *Creb5* and *Tril. Creb5* codes for cyclic AMP-Responsive Element-Binding Protein 5. *Creb5* has high expression in the heart (Fagerberg *et al.* 2014) and has been implicated in cardiac function and pathology (Schisler *et al.* 2015). *Tril* is Tlr4 interactor with leucine-rich repeats and is a functional component of Tlr4 complex involved with LPS signaling and is highly expressed in the kidney (Carpenter *et al.* 2009), indicating a potential role for *Tril* in blood pressure regulation. *Tril*(-/-) mice also produce lower levels of multiple proinflammatory cytokines and chemokines within the brain after E. coli and LPS challenge (Wochal *et al.* 2014), suggesting a potential role in immune modulation. There were no significant hits in the *Hhs2* locus.

Further experimental validation will be required to confirm the association between our any of the above candidates and Hhs. However, the above genes each have compelling functional associations that can inform follow up studies.

### Computation and quantitative trait gene prediction

Definitive functional validation of a quantitative trait gene (QTG) has traditionally required either congenic mapping to resolve an extremely narrow QTL, or *ad hoc* nomination of a candidate gene for direct experimentation. The advent of modern genetic technologies, such as CRISPR/Cas9 (Hsu *et al.* 2014), allow relatively fast and inexpensive allelic manipulations, so the burden of QTG prediction is moving toward a regime in which a small handful of strong candidates can be followed up individually. Importantly, many QTLs, including *Hhs*, contain multiple causal variants, so fine-mapping alone cannot provide definitive validation. Therefore, computational tools that can identify a small number of reasonable candidates can be a significant aid in biological follow-up. We have presented an integrative strategy for ranking genes in a QTL by combining predicted functional associations to the trait with SNP associations. Our method produces a full ranked list of genes in the locus providing researchers with the potential to validate multiple targets. To this end, the *Hhs* QTL represents an extreme use case for QTG prediction–a large, polygenic QTL associated with a physiologically complex trait.

One major limitation to our approach is the decision of which functional terms to include for network-based prediction. The better tailored this set is to the trait of interest, the greater confidence we can have in the final predictions. In principle, the inclusion of a spurious functional term could skew the rankings toward genes that are functionally associated with the spurious term but irrelevant to the trait of interest. One potential way around this issue is to use functional data, such as transcriptomics, directly from the mapping population. However, in some cases, including Hhs, the relevant tissue in which to measure gene expression may not be obvious. Alternatively, one could consider distinct rankings for each functional term. In any case, the researcher will have to exercise some measure of judgment in the prioritization process. However, by transferring the judgments from a large list of positional candidate genes to a smaller and more tractable list of trait-related biological processes, we have shown that we can arrive at a strong set of follow up candidates that would have evaded naive *p* value filters and are relatively unbiased by findings published in the literature.

The final output of our method, a ranked list of positional candidate genes, is easy to interpret, and provides researchers with a clear set of hypotheses to test in the lab. While this approach cannot definitively identify the causal gene or genes in a locus, it does provide a much-reduced set of plausible candidates to test.

## DATA AVAILABILITY

A reproducible workflow in R markdown is available on GitHub (https://github.com/MahoneyLab/HhsFunctionalRankings). This workflow contains all code required to reproduce the figures and results presented in this manuscript.

The data used as input for the workflow, as well as intermediate and final results are available on Figshare (https://figshare.com).

## ACKNOWLEDGMENTS

We would like to thank Laura Cort for supervising students during genotyping of congenic mice. ALT and JMM are supported by a grant (R21 LM012615) from the National Library of Medicine of the United States National Institutes of Health (NIH). AR, DNK, EPB, and CT were supported by grants from the NIH and the National Multiple Sclerosis Society (NMSS). DNK was supported by NIH grants from the National Institute of Neurological Disease and Stroke (R01 NS097596), National Institute of Allergy and Infectious Disease (R21 AI145306), and the NMSS (RR-1602-07780).

## Supplemental Files

**Supplemental File 1 Table of PCR primers for genotyping**. An Excel file listing the primers for genotyping microsattelite markers.

**Supplemental File 2 Table of gene module enrichments**. A tab-delimited file listing the top enrichment terms for each module. Columns are Term: the term name, Module: the number of the module within the term, N.Genes: the number of genes in the module, Enrichment.Terms: the significantly (*p* ≤ 0.05) enriched terms associated with the genes in the module.

**Supplemental File 3 A tab-delimited table listing SNPs that were assigned to genes**. Each SNP was assigned to the nearest gene within 1Mb. The table contains six columns: SNP (the rs number of each SNP), Chr (the chromosome on which the SNP is located), Position (the genomic position in bp of each SNP), Nearest.Gene (the nearest protein coding gene), Distance_to_gene (the distance in bp to the listed gene), p.value (the *p*_*EMMA*_ of each SNP).

**Supplemental File 4 Tab-delimited table containing the** *p*_*EMMA*_ **for all SNPs in the *Hhs* locus**. The table contains four columns: refsnp_id (the rs number for each SNP), chr_name (the chromosome each SNP is found on, chrom_start (the genomic position of eacn SNP in bp), p.value (*p*_*EMMA*_).

**Supplemental File 5 Final gene scores (***S*_*cg*_**) for genes in the *Hhs* locus**. Tab-delimited table with four columns: gene.name (name of ranked gene), gene.position (genomic location of the gene in bp), EMMa.p (*p*_*EMMA*_), FP (false positive rate of SVM score), gene.final.score (the sum of *p*_*EMMA*_ and FP).

**Supplemental File 6 Cycling conditions for PCR**. A .docx file containing cycling conditions for SNP genotyping.

**Supplemental File 7 Gene lists used for training**. Zipped file containing all gene lists used in the analysis.

**Supplemental File 8 Collated genotypes across *Hhs* locus for 23 inbred strains**. A comma-separated table indicating the genotypes of 23 inbred strains for 13,598 SNPs in the *Hhs* locus (45-128 Mbp). There are 28 columns in this table: chr (the chromosome the SNP is located on), bp38 (the genomic coordinates of the SNP in bp, genome build 38 (mm10)), rs (SNP rs number), observed (genotypes observed at the locus), dbsnp142annot (gene annotation of SNP), and the genotypes for each of the 23 strains.

## LITERATURE CITED

Ashburner, M., C. A. Ball, J. A. Blake, D. Botstein, H. Butler, et al., 2000 Gene ontology: tool for the unification of biology. Nature Genetics 25: 25.

Baker, E. J., J. J. Jay, J. A. Bubier, M. A. Langston, and E. J. Chesler, 2012 GeneWeaver: a web-based system for integrative functional genomics. Nucleic Acids Research 40: D1067–76.

Barrett, T., S. E. Wilhite, P. Ledoux, C. Evangelista, I. F. Kim, et al., 2012 NCBI GEO: archive for functional genomics data sets—update. Nucleic Acids Research 41: D991–D995.

Benson, M. D., C. C. Khor, P. J. Gage, and O. J. Lehmann, 2017 A targeted approach to genome-wide studies reveals new genetic associations with central corneal thickness. Molecular Vision 23: 952.

Bunce, M., C. O’neill, M. Barnardo, P. Krausa, M. Browning, et al., 1995 Phototyping: comprehensive dna typing for hla-a, b, c, drb1, drb3, drb4, drb5 & dqb1 by pcr with 144 primer mixes utilizing sequence-specific primers (pcr-ssp). Tissue antigens 46: 355–367.

Carpenter, S., T. Carlson, J. Dellacasagrande, A. Garcia, S. Gibbons, et al., 2009 TRIL, a functional component of the TLR4 signaling complex, highly expressed in brain. The Journal of Immunology 183: 3989–3995.

Csardi, G., 2006 The igraph software package for complex network research. InterJournal, Complex Systems p. 1695.

Durinck, S., Y. Moreau, A. Kasprzyk, S. Davis, B. De Moor, et al., 2005 BioMart and Bioconductor: a powerful link between biological databases and microarray data analysis. Bioinformatics 21: 3439–3440.

Durinck, S., P. T. Spellman, E. Birney, and W. Huber, 2009 Mapping identifiers for the integration of genomic datasets with the R/Bioconductor package biomaRt. Nature Protocols 4: 1184.

Edgar, R., M. Domrachev, and A. E. Lash, 2002 Gene Expression Omnibus: NCBI gene expression and hybridization array data repository. Nucleic Acids Research 30: 207–210.

Elmore, J. R., M. A. Obmann, H. Kuivaniemi, G. Tromp, G. S. Gerhard, et al., 2009 Identification of a genetic variant associated with abdominal aortic aneurysms on chromosome 3p12.3 by genome wide association. Journal of Vascular Surgery 49: 1525–1531.

Fagerberg, L., B. M. Hallström, P. Oksvold, C. Kampf, D. Djureinovic, et al., 2014 Analysis of the human tissue-specific expression by genome-wide integration of transcriptomics and antibody-based proteomics. Molecular & Cellular Proteomics 13: 397–406.

Gene Ontology Consortium, 2018 The Gene Ontology resource: 20 years and still GOing strong. Nucleic Acids Research 47: D330–D338.

Ghannadan, M., A. W. Hauswirth, G.-H. Schernthaner, M. R. Müller, W. Klepetko, et al., 2002 Detection of novel CD antigens on the surface of human mast cells and basophils. International Archives of Allergy and Immunology 127: 299–307.

Goya, J., A. K. Wong, V. Yao, A. Krishnan, M. Homilius, et al., 2015 FNTM: a server for predicting functional networks of tissues in mouse. Nucleic Acids Research 43: W182–7.

Greene, C. S., A. Krishnan, A. K. Wong, E. Ricciotti, R. A. Zelaya, et al., 2015 Understanding multicellular function and disease with human tissue-specific networks. Nature Genetics 47: 569–576.

Guan, Y., C. L. Ackert-Bicknell, B. Kell, O. G. Troyanskaya, and M. A. Hibbs, 2010 Functional genomics complements quantitative genetics in identifying disease-gene associations. PLoS Computational Biology 6: e1000991.

Hedley, P. L., P. Jørgensen, S. Schlamowitz, J. Moolman-Smook, J. K. Kanters, et al., 2009 The genetic basis of Brugada syndrome: a mutation update. Human Mutation 30: 1256–1266.

Hill, S., C. Ganellin, H. Timmerman, J. Schwartz, N. Shankley, et al., 1997 International union of pharmacology. XIII. classification of histamine receptors. Pharmacological Reviews 49: 253–278.

Hiltunen, J. O., A. Laurikainen, M. S. Airaksinen, and M. Saarma, 2000 GDNF family receptors in the embryonic and postnatal rat heart and reduced cholinergic innervation in mice hearts lacking RET or GFR*α*2. Developmental Dynamics 219: 28–39.

Hsu, P. D., E. S. Lander, and F. Zhang, 2014 Development and applications of CRISPR-Cas9 for genome engineering. Cell 157: 1262–1278.

Ju, W., C. S. Greene, F. Eichinger, V. Nair, J. B. Hodgin, et al., 2013 Defining cell-type specificity at the transcriptional level in human disease. Genome Research 23: 1862–1873.

Kang, H. M., N. A. Zaitlen, C. M. Wade, A. Kirby, D. Heckerman, et al., 2008 Efficient control of population structure in model organism association mapping. Genetics 178: 1709–1723.

Krishnan, A., R. Zhang, V. Yao, C. L. Theesfeld, A. K. Wong, et al., 2016 Genome-wide prediction and functional characterization of the genetic basis of autism spectrum disorder. Nature Neuroscience 19: 1454.

Lieberman, P. and L. H. Garvey, 2016 Mast cells and anaphylaxis. Current Allergy and Asthma Reports 16: 20.

Meyer, D., E. Dimitriadou, K. Hornik, A. Weingessel, and F. Leisch, 2018 e1071: Misc Functions of the Department of Statistics, Probability Theory Group (Formerly: E1071), TU Wien. R package version 1.7-0.

Motenko, H., S. B. Neuhauser, M. O’keefe, and J. E. Richardson, 2015 MouseMine: a new data warehouse for MGI. Mammalian Genome 26: 325–330.

Mouse Genome Informatics, Mouse Genome Informatics Web Site http://www.informatics.jax.org/. Accessed May 23, 2019.

Napolitano, C., I. Splawski, K. W. Timothy, R. Bloise, and S. G. Priori, 2015 Timothy syndrome. In GeneReviews, University of Washington, Seattle.

Newman, M. E. J., 2004 Fast algorithm for detecting community structure in networks. Physical Review E 69: 066133.

Parker, C. C., G. Sokoloff, E. Leung, S. L. Kirkpatrick, and A. A. Palmer, 2013 A large QTL for fear and anxiety mapped using an F2 cross can be dissected into multiple smaller QTLs. Genes, Brain and Behavior 12: 714–722.

Petkov, P. M., Y. Ding, M. A. Cassell, W. Zhang, G. Wagner, et al., 2004 An efficient snp system for mouse genome scanning and elucidating strain relationships. Genome research 14: 1806–1811.

Raza, A., E. C. Chan, W.-S. Chen, Z. Xie, L. M. Scott, et al., Under Review A natural mouse model reveals genetic determinants of Systemic Capillary Leak Syndrome (Clarkson disease).

Reimand, J., R. Kolde, and T. Arak, 2018 gProfileR: Interface to the ‘g:Profiler’ Toolkit. R package version 0.6.7.

Rusmini, M., P. Griseri, F. Lantieri, I. Matera, K. L. Hudspeth, et al., 2013 Induction of RET dependent and independent proinflammatory programs in human peripheral blood mononuclear cells from Hirschsprung patients. PLoS One 8: e59066.

Schisler, J. C., T. J. Grevengoed, F. Pascual, D. E. Cooper, J. M. Ellis, et al., 2015 Cardiac energy dependence on glucose increases metabolites related to glutathione and activates metabolic genes controlled by mechanistic target of rapamycin. Journal of the American Heart Association 4: e001136.

Smith, C. L. and J. T. Eppig, 2012 The Mammalian Phenotype Ontology as a unifying standard for experimental and highthroughput phenotyping data. Mammalian Genome 23: 653–668.

Sudweeks, J. D., J. A. Todd, E. P. Blankenhorn, B. B. Wardell, S. R. Woodward, et al., 1993 Locus controlling bordetella pertussisinduced histamine sensitization (bphs), an autoimmune diseasesusceptibility gene, maps distal to t-cell receptor beta-chain gene on mouse chromosome 6. Proceedings of the National Academy of Sciences 90: 3700–3704.

Vaz, N. M., C. M. de Souza, M. M. Hornbrook, D. G. Hanson, and N. R. Lynch, 1977 Sensitivity to intravenous injections of histamine and serotonin in inbred mouse strains. International Archives of Allergy and Immunology 53: 545–554.

Wang, M., T. Shibamoto, M. Tanida, Y. Kuda, and Y. Kurata, 2014 Mouse anaphylactic shock is caused by reduced cardiac output, but not by systemic vasodilatation or pulmonary vasoconstriction, via paf and histamine. Life Sciences 116: 98–105.

Wochal, P., V. A. Rathinam, A. Dunne, T. Carlson, W. Kuang, et al., 2014 TRIL is involved in cytokine production in the brain following *escherichia coli* infection. The Journal of Immunology 193: 1911–1919.

Xie, Z., V. Nagarajan, D. E. Sturdevant, S. Iwaki, E. Chan, et al., 2013 Genome-wide SNP analysis of the systemic capillary leak syndrome (clarkson disease). Rare Diseases 1: e27445.

Yan, J., Q. Li, Y. Luo, S. Yan, Y. He, et al., 2018 Association of CACNA1C gene genetic polymorphism with the susceptibility as well as prognosis for chronic spontaneous urticaria. Zhong nan da xue xue bao. Yi xue ban= Journal of Central South University. Medical sciences 43: 929–936.

Yazbek, S. N., D. A. Buchner, J. M. Geisinger, L. C. Burrage, S. H. Spiezio, et al., 2011 Deep congenic analysis identifies many strong, context-dependent QTLs, one of which, *Slc35b4*, regulates obesity and glucose homeostasis. Genome Research 21: 1065–1073.

